# Range-wide validation of reduced locomotor endurance in unisexual *Ambystoma* salamanders

**DOI:** 10.64898/2026.05.29.728744

**Authors:** Bethanie Majewski, Jessica Castetter, Gabriela Bilbrey, Robert D. Denton

## Abstract

Locomotor endurance is a critical physiological trait dictating terrestrial dispersal and metapopulation connectivity, especially in amphibians. The unisexual *Ambystoma* complex is an ancient, all-female polyploid lineage that reproduces via kleptogenesis. This unique reproductive mode creates an evolutionary mismatch between a conserved mitochondrial genome and divergent nuclear subgenomes that are taken from sympatric, sexual species. This provides a compelling system for testing the physiological limits of polyploidy and how mitonuclear dissonance and subgenome composition influence phenotypes. Previous locomotor assessments of this lineage demonstrated that polyploid salamanders display reduced locomotor endurance compared to sexual species. To overcome previous limitations in geographic sampling and sample size, we conducted standardized treadmill endurance trials on a broad geographic sampling of 102 salamanders, comparing the performance of triploid unisexual biotypes (LJJ and LLJ) directly to their sexually reproducing parental species (*A. jeffersonianum* and *A. laterale*). We found that reproductive mode significantly dictated endurance performance. Sexual species broadly outperformed unisexuals, driven by *A. jeffersonianum* exceeding all unisexual biotypes and *A. laterale* outperforming LJJ unisexuals. Within the unisexual cohort, subgenome dosage did not influence performance: LJJ individuals and LLJ individuals had similar performance.. We propose that the aerobic capacity of unisexual salamanders is limited, potentially due to mitonuclear mismatch or the biophysical constraints of increased cellular volume, but not due to the contributions of individual subgenomes. This endurance deficit is likely to restrict unisexual salamanders’ dispersal capabilities, leaving populations uniquely vulnerable to ongoing habitat fragmentation compared to more mobile, sexual species while potentially preventing unisexuals from driving sexual hosts locally extinct.

## Introduction

Across the animal kingdom, locomotor endurance serves as a fundamental performance trait that is crucial for overall survival and reproductive success, dictating an organism’s capacity to engage in mate searching, active foraging, predator evasion, and dispersal (Husak and Lailvaux, 2019; Llewelyn et al., 2010; Ramos et al., 2004; Seebacher and Walter, 2012). But prolonged physical exertion requires significant energy, making it one of the most primary aspects of an organism’s energy allocation strategy and a focal point in contemporary evolutionary ecophysiology (Husak et al., 2016). Within populations that must routinely move between distinct habitats, locomotor endurance is the central physiological mechanism enabling successful migration and population connectivity (Alerstam and Bäckman, 2018; Bowler and Benton, 2005). Amphibians epitomize this life-history challenge, as the majority of pond-breeding species undergo obligate seasonal migrations between terrestrial overwintering habitats and aquatic vernal pools for reproduction (Petranka, 1998). Amphibian survival during these terrestrial migrations is severely constrained by physiological constraints, primarily the necessity to maintain critical hydration levels and regulate body temperature (Cayuela et al., 2020; Galindo et al., 2023; Hillman et al., 2014; Roznik et al., 2018). At the landscape scale, dispersal drives metapopulation dynamics, preventing local extinction events through the continuous influx of genetic diversity across fragmented populations (Marsh and Trenham, 2001; Smith and Green, 2005).

The capability of an amphibian to successfully disperse across a landscape is governed by both extrinsic factors (e.g. habitat patch size, inter-patch distance, climatic fluctuations, social cues, competition, etc.) and intrinsic organismal traits, including body size, sex, and baseline metabolic rates (Bailey et al., 2018; Bredeweg et al., 2019; Le Galliard et al., 2004; Schuler et al., 2017; Teunissen et al., 2025). For these intrinsic factors, parameters such as oxygen consumption rates, muscle isometric tension, and thermal acclimation significantly dictate the limits of locomotor capacity in salamander populations (Bennett et al., 1989; Finkler et al., 2003; Johnson et al., 2010). At the core of this metabolic output are mitochondria, which drive aerobic energy production. Consequently, variations in mitochondrial efficiency exert profound downstream effects on organismal physiology, behavioral ecology, and sustained locomotor endurance (Anderson et al., 2022). However, cellular respiration does not rely on the mitochondrial genome alone. It requires coordination between proteins encoded by both mitochondrial DNA (mtDNA) and nuclear DNA (nuDNA). When evolutionarily divergent mtDNA and nuDNA are present in the same cells, the resulting mitonuclear dissonance can disrupt electron transport chain efficiency, leading to reductions in physiological capacity (Anderson et al., 2022; Burger et al., 2025; Klabacka et al., 2022; Koch et al., 2025; Moran et al., 2024; Nagao et al., 1998; Rodríguez et al., 2021). However, the connection between mitonuclear dissonance and locomotor endurance is relatively unexplored.

In this study, we test the hypothesis that amphibians with inherent evolutionary dissonance between their mitochondrial and nuclear genomes will exhibit diminished locomotor endurance compared to species possessing strictly co-evolved mtDNA and nuDNA. To test this hypothesis, we use the unique, unisexual (all-female) salamander complex within the genus *Ambystoma*. Unisexual *Ambystoma* provide a unique study system for this experiment: they are the oldest known lineage of unisexual vertebrates (mtDNA lineage is ~5 million years old) but continually acquire new nuDNA from sympatric sexual species (Bi and Bogart, 2006; Bi and Bogart, 2010; Bogart et al., 2007). This unique reproductive system–called kleptogenesis–requires unisexual females to use sperm from sympatric sexual males to trigger egg development. While the resulting offspring are usually clonal, the females occasionally incorporate the male’s nuclear DNA via ploidy elevation or genome replacement (Bi et al., 2008; Bogart, 2019; Bogart et al., 2007). Over millions of years, this has resulted in a monophyletic, highly conserved mitochondrial lineage paired with vastly divergent nuclear genomes (Hedges et al., 1992). Because kleptogenesis forces an ancient mitochondrial genome to interact with a constantly shifting array of nuclear subgenomes, unisexual *Ambystoma* would theoretically show perpetual mitonuclear dissonance as new nuDNA is swapped into a population. Previous empirical work has demonstrated that the physiological costs of this mismatch can manifest as significantly reduced locomotor endurance and shorter realized dispersal distances in field populations (Denton et al., 2017). However, polyploidy is also likely to play a role in these comparisons between unisexual salamanders and their sexual relatives, but there are few examples of systems were the effects of polyploidy and mitonuclear dissonance are readily disentangled (Greimann et al., 2020 as a potential example). Recent multi-trait analyses further confirm that the specific genomic composition of these hybrids serves as a direct predictor of their physiological performance, highlighting deep physiological trade-offs between traits such as resistance to water loss and carbon dioxide respiration rates (Burger et al., 2025).

Despite clear differences in functional physiology between unisexual and sexual salamanders, previous locomotor experiments have been limited in scope, focusing on restricted geographic ranges and small sample sizes (Denton et al., 2017). As in many other pond breeding amphibians, movements of *Ambystoma* salamanders are primarily constrained to breeding season migrations to and from nearby wetlands or rare dispersal events to new wetland sites (Petranka, 1998). Here, we apply a standardized, rigorous treadmill endurance methodology to a comprehensive sample of salamanders from across the entire geographic range of the *Ambystoma* complex. We investigate whether unisexual salamanders broadly exhibit reduced locomotor capabilities, and whether specific combinations of subgenomes consistently predict their physiological performance limits. We tested two specific hypotheses. First, we hypothesized that the mitonuclear coevolution within sexually reproducing salamander species (*A. laterale* and *A. jeffersonianum*) would allow for greater locomotor endurance. We predicted that these sexual species would have significantly greater total distance traveled before exhaustion compared to unisexual salamanders with putative mitonuclear dissonance. More specifically, we expected this performance gap to be most pronounced when comparing unisexuals to *A. jeffersonianum*, given the latter’s larger body size and relationship with more heterogeneous terrain. Second, we hypothesized that within the unisexual salamanders, distinct polyploid biotypes would influence locomotor endurance. We predicted that unisexual individuals would perform most closely to the sexual species with which they share the greatest number of subgenomes. Specifically, the triploid LLJ (two *A. laterale* genomes, one *A. jeffersonianum* genome) and LJJ (one *A. laterale* genome, two *A. jeffersonianum* genomes) combinations, would exhibit locomotor performance more like *A. laterale* and *A. jeffersonianum*, respectively.

## Materials and methods

We tested 102 adult salamanders (unisexual LJJ, unisexual LLJ, *A. jeffersonianum*, *A. laterale*) that were collected from across their ranges between 2019-2023 according to local state wildlife regulations and housed on the authors’ institution (Table 1). All salamanders had been in captivity for at least one year at the time of this experiment, during which time they were housed individually inside plastic bins within a climate-controlled refrigerator (~13°C; IACUC #RD2021-2). Both sexual species were a mix of male and female individuals. As unisexual individuals cannot be reliably identified morphologically, we conducted a genetic screening procedure to A) confirm the species assignment of each animal and B) infer the chromosome composition of identified unisexuals. First, we sequenced a high variable portion of the mitochondrial genome (primers F-THR and R-651; McKnight and Shaffer, 1997) and assigned individuals to either the unisexual lineage, *A. laterale*, or *A. jeffersonianum* using reference sequences. For individuals assigned into the unisexual lineage, we genotyped them at three microsatellite loci that differentially amplify in each sexual species (Teltser and Greenwald, 2015). This process, along with *a priori* expectations based on species ranges, has been used to successfully characterize the biotype (e.g. LLJ, LJJ, etc) in multiple studies across a broad geographic range (Bogart and Klemens, 2008; Bogart et al., 2007; Julian et al., 2003; Ramsden et al., 2006).

**Table 1.** Summary of individuals used in locomotor trials. Individuals who captured animals reported sex based on external breeding morphology at the time of capture and genetic analyses were used to separate samples into species/lineage (Ambystoma laterale, A. jeffersonianum, or unisexuals) and then biotype for the samples identified as part of the unisexual lineage.

| Location (County, State) | Number of individuals (male/female/unknown) |  |  |  |
| --- | --- | --- | --- | --- |
|  | Unisexual (LLJ) | Unisexual (LJJ) | *Ambystoma jeffersonianum* | *Ambystoma laterale* |
| Litchfield, CT | - | 0/5/0 | 1/2/0 | - |
| Middlesex, CT | - | - | - | 0/1/0 |
| Windham, CT | - | - | - | 1/3/0 |
| Campbell, KY | - | 0/7/0 | 7/0/0 | - |
| Jessamine, KY | - | - | 0/1/0 | - |
| Kenton, KY | - | 0/4/0 | - | - |
| Waldo, ME | 0/0/1 | - | - | - |
| Barry, MI | 0/8/0 | - | - | 0/1/0 |
| Livingston, MI | 0/10/0 | - | - | 8/1/6 |
| Mower, MN | - | - | - | 0/1/0 |
| Monroe, NY | 0/1/0 | 0/1/0 | - | 0/1/0 |
| Crawford, OH | - | 0/1/0 | 1/0/0 | - |
| Delaware, OH | - | 0/3/0 | 5/0/0 | - |
| Addison, VT | 0/0/4 | 0/2/0 | - | - |
| Calumet, WI | 0/12/0 | - | - | 2/0/0 |
| Total | 36 | 23 | 17 (14/3/0) | 25 (11/8/6) |

To quantify locomotor endurance, we conducted treadmill trials with each individual salamander. Because of the established repeatability of locomotor endurance trials across both long and short periods of time (Austin and Shaffer, 1992; Hudson et al., 2016), other studies show no difference in performance between recently captured animals and those held in captivity (Denton et al., 2017), and our desire to compare measurements to previous work, we chose to proceed with animals acclimated to captivity and each animal was only tested once to minimize potential stress. We randomly selected individuals to participate daily throughout the summer of 2024. We conducted 2-5 trials per day, depending on the availability of researchers. Schedules were determined in advance in order to keep food intake standardized (Hudson et al., 2016). We fed each animal exactly 12 days and 2 days before their trial. Each meal consisted of one appropriately-sized worm (*Eisenia hortensis*) cultured in the lab on a diet of oats, cornmeal, and calcium supplements. If an animal did not eat on their scheduled feeding day, they were moved to a date later in the summer. On the day of the trial, we weighed the salamanders and measured their snout to posterior vent (SVL) length. Before the specific trial, the salamander was brought out of the refrigerator and allowed to acclimate to room temperature (room cooled by air conditioning unit to 16-19°C) in their bin for twenty minutes.

After acclimation, we placed the salamander onto the custom treadmill, consisting of a neoprene wetsuit material within a plexiglass box that is rotated by a variable speed motor. This exact treadmill was used previously for *Ambystoma* salamanders (Denton et al., 2017; Johnson et al., 2010). When the salamander began to move, we would begin moving the tread at a pace equivalent to the salamander’s speed. When animals stopped moving, we used a metal spatula to gently prod the animal on the tail and back legs. After three minutes of walking, we stopped the treadmill and conducted a righting response test. The salamander was flipped on its back and allowed 3 seconds to right itself (Austin and Shaffer, 1992; Shaffer et al., 1991). If the salamander could right itself within 3 seconds, then we sprayed it and the tread with water via a spray bottle and allowed them to continue their trial on the treadmill. If the salamander did not successfully right itself after 3 seconds, then that specific trial was concluded. In cases where the salamander refused to walk, we ended their trial after 10 minutes. At the conclusion of each trial, we sprayed the animal with water again, returned it to their bin in the refrigerator, and monitored their behavior closely for the following three days. For each animal, we recorded the total distance traveled (m), the amount of active time on the treadmill, and the reason for ending each trial (failed righting response or timed out).

All statistical analyses and visualizations were conducted in R version 4.3 (R Core Team, 2023) using the packages lme4 (Bates et al., 2015), lmerTest (Kuznetsova et al., 2017), emmeans (Lenth, 2023), and tidyverse (Wickham et al., 2019). Prior to constructing lineage models, we evaluated collinearity between the different candidate morphological covariates. We found that salamander body mass (weight in g) and SVL (cm) were strongly collinear (Spearman’s rank correlation, *S* = 24,933, *p* < 0.001). To control for body size without collinearity bias, we calculated a Scaled Mass Index (SMI; Peig and Green, 2009). The scaling exponent (*b*_SMA_) was determined via Standard Major Axis (SMA) regression of ln(Weight) on ln(SVL), calculated as the ratio of their sample standard deviations. Each individual’s body condition was subsequently calculated using the SMI formula.

Locomotor endurance (the total distance traveled prior to failed righting response) was log_10_ transformed to achieve homoscedasticity. After visually inspecting the distribution of endurance estimates, we removed a single extreme outlier that was more than 1.5 the Interquartile Range (IQR). In addition, we removed the single individual that did not have a failed righting response and timed out its trial. Together, this reduced the final dataset from 102 individuals to 100 individuals (17 *A. jeffersonianum*, 25 *A. laterale*, 22 LJJ unisexuals, and 36 LLJ unisexuals). Before proceeding to test our hypotheses, we determined that sex had no influence on locomotor endurance (F_1,21.1_ = 0.040, *p* = 0.844) using a linear mixed-effects model restricted exclusively to the two sexual species (LogDistance as the response, Biotype, SMI, and Sex as fixed effects, and capture site as a random intercept). With no sexual dimorphism in endurance, sexes were pooled within the sexual species for all subsequent analyses.

To evaluate our hypotheses, we modeled locomotor endurance using Linear Mixed-Effects Models (LMM). We used capture sites as a random intercept in all models. First, to test the contrast between reproductive strategies broadly, we modeled LogDistance as a function of reproductive mode (sexual vs unisexual) with SMI as a continuous covariate. Second, to test for subgenome dosage effects, we modeled Log Distance as a function of biotype (JJ, LL, LJJ, LLJ) and SMI. Model assumptions were verified using diagnostic residual plots (residuals vs fitted values and normal Q-Q plots) and confirmed via the Shapiro-Wilk normality test (*W* = 0.977, *p* = 0.072). We then conducted post-hoc pairwise contrasts across biotypes on Estimated Marginal Means (EMMs) derived from the fitted mixed-effects model using Tukey’s Honest Significant Difference (HSD) adjustment for multiple comparisons. To provide biological interpretability, we back-transformed to express mean fold-differences and percentage change between groups. Finally, we evaluated whether differences in endurance could be driven by movement speed rather than stamina. We calculated locomotor speed (m/s) for each trial. Because raw speeds deviated significantly from assumptions (Shapiro-Wilk *W* = 0.126, *p* < 0.0001), we tested for speed differences between groups using a non-parametric Kruskal-Wallis test and between reproductive modes using a Mann-Whitney *<u>U</u>* <u>test.</u>

## Results

Locomotor endurance differed significantly between reproductive strategies. After accounting for capture locality and body condition, sexually reproducing salamanders exhibited higher endurance than unisexual salamanders (*F_1,19.9_* = 35.58, *p* < 0.0001; Welch’s two-sample t-test: t_88.6_ = 5.365, *p* < 0.0001; Figure 1). Individuals from sexual species traveled a mean log distance of 1.607 ± 0.057 SE (geometric mean = 40.46 m, n = 42), whereas unisexual individuals traveled a mean log distance of 1.201 ± 0.050 SE (geometric mean = 15.88 m, n = 58). When we back-transformed the model estimates, these values reveal that sexually reproducing species traveled an average of 2.55-fold farther before exhaustion compared to the unisexuals.

**Figure 1.**
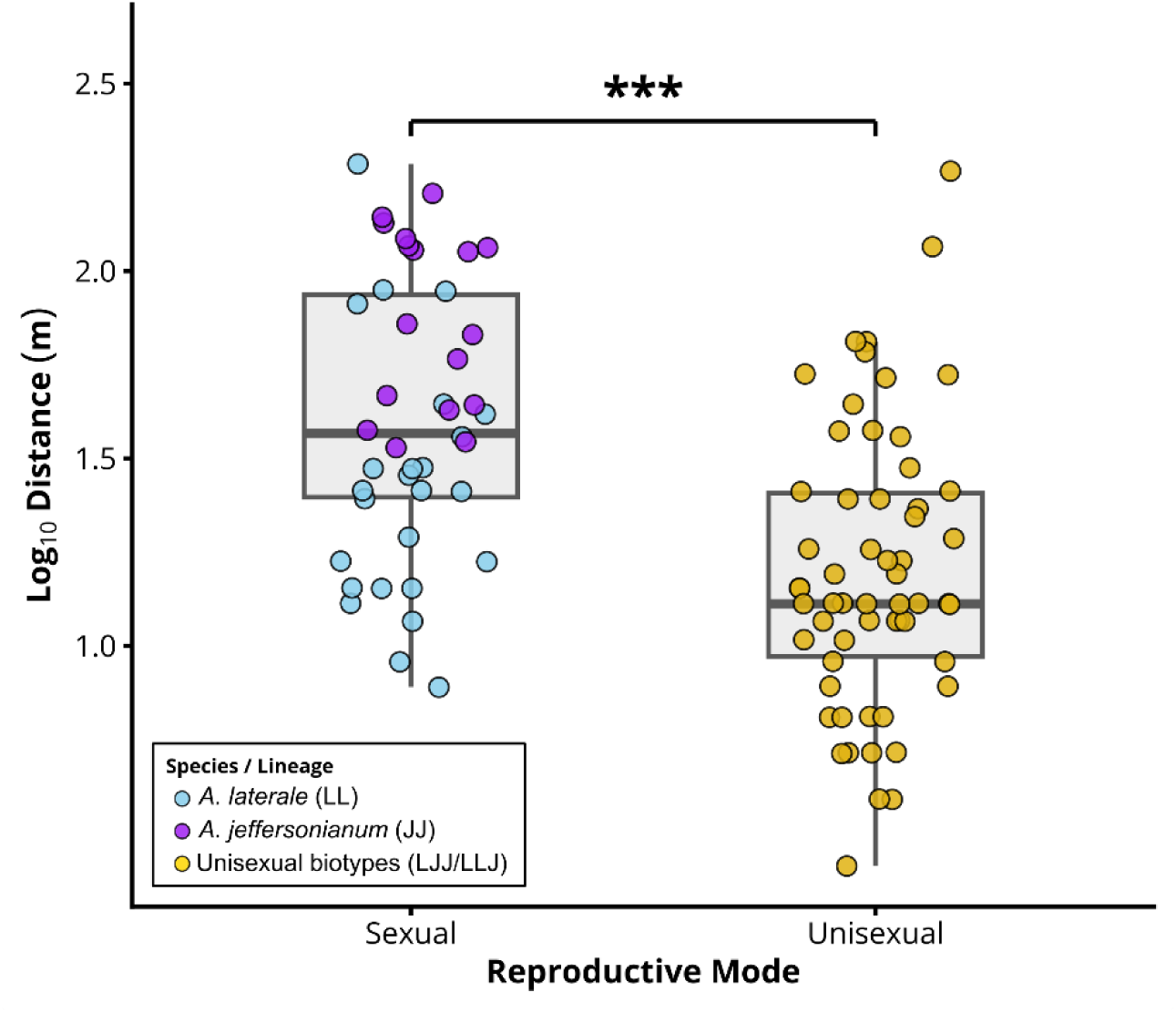
Locomotor endurance by reproductive strategy. Boxplots represent Log_10_ distance traveled by salamanders of two sexual species (Ambystoma laterale and A. jeffersonianum) and two biotypes of a unisexual lineage (LLJ or LJJ). Jittered points represent individual trials, color-coded by taxonomic identity. Sexually reproducing individuals traveled significantly farther than unisexuals before reaching a failure of righting response (Type III LMM, p < 0.001; Welch’s t-test: t_88.6_ = 5.365, p < 0.0001 denoted by ***).

For the primary model evaluating genome dosage, we found that biotype was the most prominent predictor of locomotor endurance (*F*_3,57.9_ = 16.233, *p* < 0.0001; Table 2). The SMI was not significant across the complete data set (*F*_1,93.9_ = 1.805, *p* = 0.182), indicating that endurance differences are not explained by body condition. The post-hoc contrasts on EMMs (Table 2, Figure 2A) revealed several significant differences. First, *A. jeffersonianum* (JJ) displayed the highest locomotor capacity of any group (1.868 ± 0.110 SE), significantly outperforming *A. laterale* (0.409 ± 0.134 SE; contrast: Δ = 0.568 ± 0.111 SE, t_39.4_ = 3.054, *p* = 0.020; Figure 2A). This corresponds to a 2.56-fold higher distance traveled in *A. jeffersonianum*. Second, *A. jeffersonianum* significantly outperformed both unisexual biotypes (*p* < 0.0001 for both contrasts), displaying a 3.70-fold increase over LJJ (Δ = 0.568 ± 0.111 SE, t_92.8_ = 5.114) and a 6.01-fold increase over LLJ (Δ = 0.779 ± 0.126 SE, t_35.7_ = 6.163). While *A. laterale* also significantly outperformed LLJ unisexuals (Δ = 0.370 ± 0.096 SE, t_84.8_ = 3.854, *p =* 0.0013, a 2.34-fold increase), there was no difference between *A. laterale* and LJJ unisexuals (Δ = 0.159 ± 0.122 SE, t_39.8_ = 1.301, *p* = 0.5679). Finally, we found no difference between unisexual biotypes (LJJ vs LLJ: Δ = 0.211 ± 0.113 SE, t_37.2_ = 1.865, *p* = 0.2605; Figure 2A).

**Figure 2.**
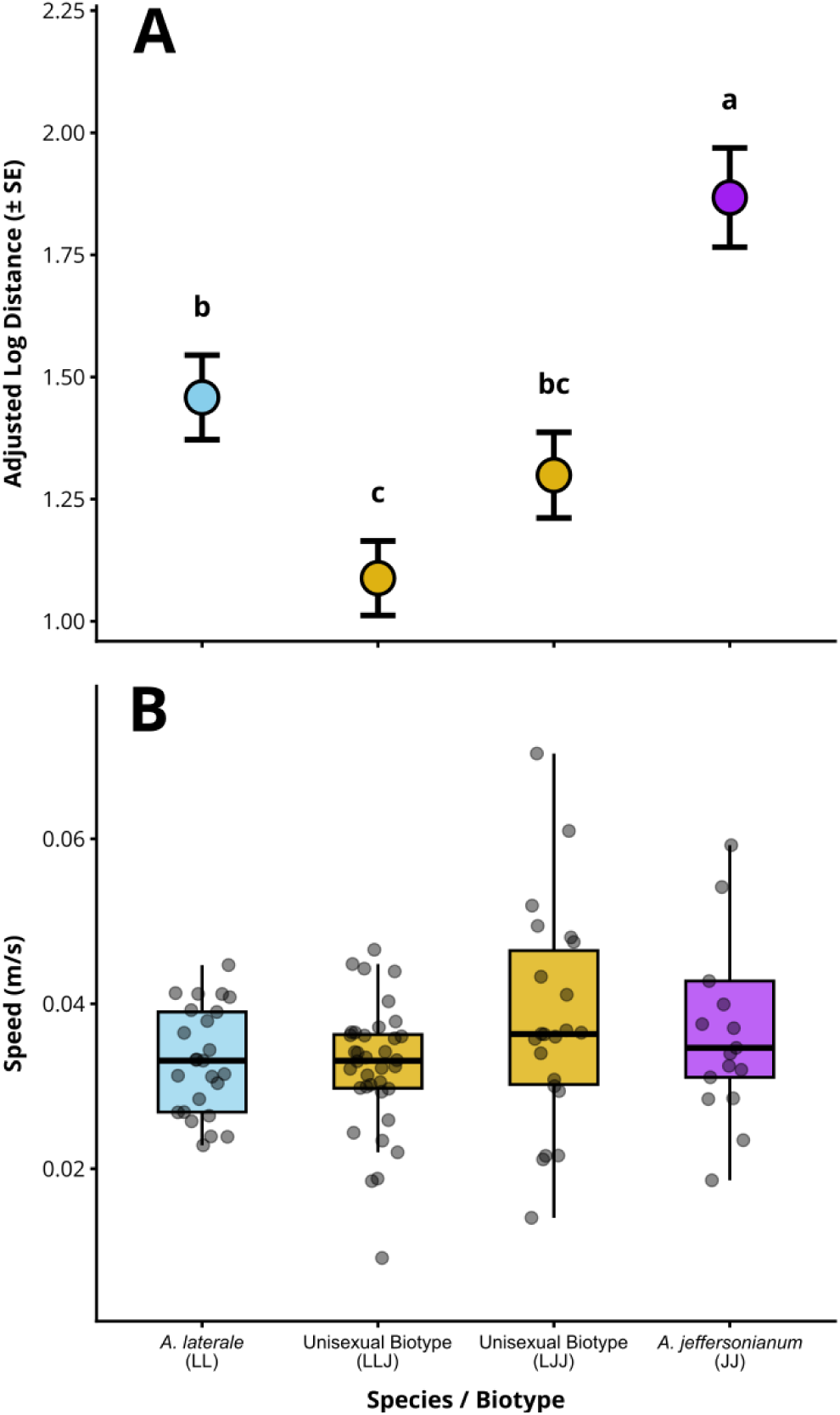
Model-adjusted locomotor endurance and speed across groups. Central points and error bars in (A) represent Estimated Marginal Means (EMMs ± SE) derived from the linear mixed-effects model accounting for scaled mass index (SMI) and capture locality. Colors represent the two sexual species (Ambystoma laterale and A. jeffersonianum) and the unisexual lineage (LLJ and LJJ biotypes). Letters above indicate statistical grouping based on Tukey-adjusted pairwise contrasts. Subgenome dosage was not supported, as LJJ and LLJ biotypes did not significantly differ (p = 0.261). Boxplots (B) summarize trial speeds (m/s) for each group along with individual values (circles). Speed was indistinguishable across groups (Kruskal-Wallis: H = 3.822, df = 3, p = 0.281).

**Table 2.** Statistical comparisons of locomotor endurance by salamanders from two species (Ambystoma laterale, A. jeffersonianum and two unisexual biotypes (LLJ unisexuals and LJJ unisexuals) using linear mixed-effects modeling of log-transformed distance. The model includes scaled mass index (SMI) as a covariate and collection site as a random intercept. Pairwise post-hoc comparisons were evaluated on Estimated Marginal Means (EMMs) with Tukey-adjusted p-values. Individual results are bolded with a p value <0.05.

| *Type III Analysis of Variance Table* |  |  |  |  |  |
| --- | --- | --- | --- | --- | --- |
| Factor | NumDF/DenDF | Sum of Squares | Mean Square | F-value | p-value |
| **Species/Biotype** | **3/57.92** | **4.828** | **1.609** | **16.233** | **<0.0001** |
| SMI | 1/93.94 | 0.179 | 0.179 | 1.805 | 0.1823 |
| *Tukey's Post-Hoc Contrasts* |  |  |  |  |  |
| Contrast | df | Estimate ± SE | t-ratio | Adjusted p | Fold Diff. (10Δ) |
| **JJ – LJJ** | **92.8** | **0.568 ± 0.111** | **5.114** | **< 0.0001** | **3.70x** |
| **JJ – LL** | **39.4** | **0.409 ± 0.134** | **3.054** | **0.0202** | **2.56x** |
| **JJ – LLJ** | **35.7** | **0.779 ± 0.126** | **6.163** | **< 0.0001** | **6.01x** |
| LJJ – LL | 39.8 | –0.159 ± 0.122 | –1.301 | 0.5679 | 0.69x |
| LJJ – LLJ | 37.2 | 0.211 ± 0.113 | 1.865 | 0.2605 | 1.63x |
| **LL – LLJ** | **84.8** | **0.37 ± 0.096** | **3.854** | **0.0013** | **2.34x** |

We found that speed was uniform across groups and could not account for disparities in distance traveled (Figure 2B). Median speed was tightly clustered between 0.033-0.036 m/s. A Kruskal-Wallis test confirmed no significant difference among the four groups (*H* = 3.822, df = 3, *p* = 0.2814). Similarly, speed did not differ between sexual and unisexual reproductive modes (Mann-Whitney *U* = 1,242.0, *p* = 0.8696).

## Discussion

The results of this study reinforce and expand the observation that sexually reproducing *Ambystoma* species possess significantly greater locomotor endurance than their sympatric unisexual counterparts (Denton et al., 2017; Figure 3). By sampling populations across diverse geographic regions, we provide confidence that the endurance deficit is a generalized, species-wide phenomenon inherent to the unisexual lineage. Sexually reproducing individuals traveled on average 2.55-fold farther before exhaustion than unisexuals after controlling for body condition and geographic origin. Furthermore, walking speed was identical across all groups, indicating that the difference in locomotor endurance reflects differences in physiological stamina and time to exhaustion rather than differences in gait, pace, or behavior.

**Figure 3.**
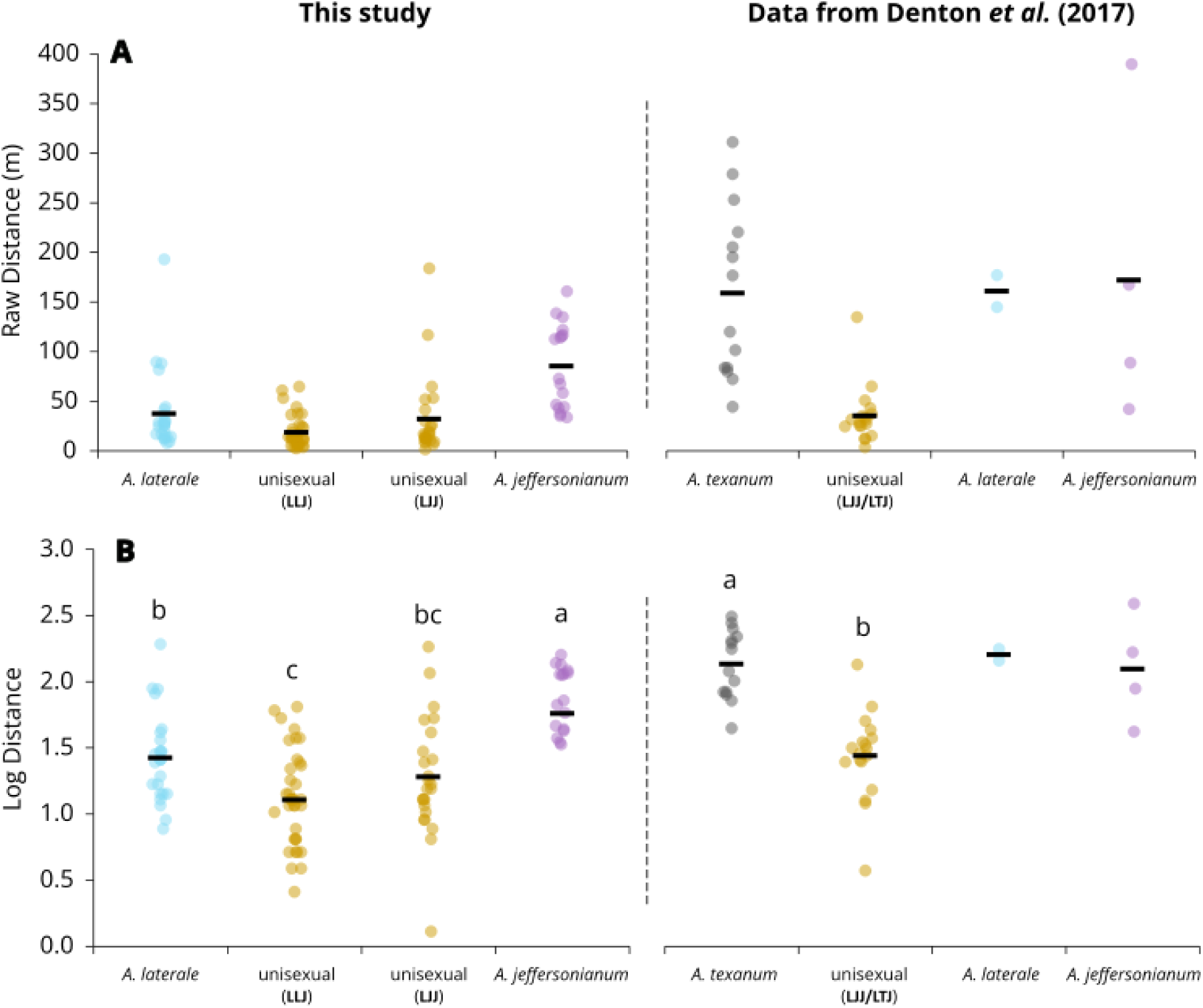
Comparison of locomotor endurance between sexual and unisexual salamanders across studies. We present both the raw distance traveled by each individual (A) and the transformed distance used in statistical analyses (B). Each colored dot is an individual animal’s trial value and horizontal black bars indicate the raw mean for each group. On the right, we report the same values from previous research (Denton et al., 2017) for comparison. Letters above groups in (B) signify statistical differences identified in each study.

Explaining the underlying mechanism for limited locomotor endurance in unisexual *Ambystoma* is difficult, as they face a complex suite of genomic and biophysical challenges that may severely restrict their endurance. They possess polyploid nuclear DNA composed of subgenomes from multiple sexual species, primarily *A. jeffersonianum* and *A. laterale*, paired with a distinct, five-million-year-old mitochondrial haplotype derived from an ancient *A. barbouri* ancestor (Bi and Bogart, 2010). Because efficient cellular respiration relies upon the structural integration of essential protein complexes encoded by both the nuclear and mitochondrial genomes, an evolutionary dissonance between these genomes can disrupt metabolic performance (Carnegie et al., 2021; Hill et al., 2019; McDiarmid et al., 2024; Rank et al., 2020). Because unisexuals, regardless of their biotype, demonstrated locomotor endurance limitations compared to at least three sexual Ambystomatids, we hypothesize that mitonuclear dissonance is the most likely mechanism that drives this functional trait.

Beyond interactions between mtDNA- and nuDNA-coded gene products, unisexual salamanders must contend with the fundamental physical constraints of polyploidy. The incorporation of two to five haploid genomes proportionately increases the absolute cell size (Austin and Bogart, 1982; Cadart et al., 2023). Because a significant portion of basal metabolic energy is dedicated to maintaining the plasma membrane, the reduction in total membrane surface area in polyploids can lower their mass-specific metabolic rate (Cadart et al., 2023). However, unisexual *Ambystoma* do not demonstrate lower whole-animal metabolic rates compared to sexual species (Burger et al., 2025). Salamanders as a group have the lowest metabolic rates among tetrapods (Chong and Mueller, 2013; Gatten et al., 1992), suggesting that the types of functional differences we observed in this study are only apparent when these species are under significant metabolic demand. When confronted with the extreme, sustained energy demands of continuous treadmill locomotion, the unisexual’s depressed metabolic ceiling may be rapidly overwhelmed, leading to more rapid muscular exhaustion. The application of this scenario to the wild is debatable, but the nature of early spring migrations and small timeframes for breeding activity are likely to emphasize rapid, sustained movements over the course of 1-3 days (Petranka, 1998). While there remain no clear empirical links between the effects of increased ploidy and locomotor endurance, it is reasonable to not consider the consequences of ploidy and mitonuclear function to be mutually exclusive. Mitonuclear dissonance and polyploid cells are likely to operate in tandem to establish a lower aerobic ceiling in unisexuals.

Within the unisexual salamanders, the two different biotypes (LLJ or LJJ) did not differ in their locomotor endurance. This result contrasts with previous studies that have demonstrated that polyploid biotypes tend to drift toward the physiological baseline of their dominant parental genome (Burger et al., 2025; Lowcock, 1994). Our results are not aligned with this genomic dosage framework: LJJ individuals (possessing two *jeffersonianum* genomes) exhibited similar locomotor endurance to LLJ individuals (possessing two *laterale* genomes). While subgenome dosage predicts static physiological traits such as thermal maximum and cutaneous water loss (Burger et al., 2025), dynamic whole-organism endurance appears constrained by a non-additive threshold. Adding a second *A. jeffersonianum* genome failed to rescue performance, broadly supporting the cause of reduced performance in unisexuals to be due to some combination of mitonuclear dissonance and ploidy rather than the individual contributions of parental genomes. To begin to disentangle these causes, future work on the unisexual salamander system should include direct measurements of mitochondrial function and sampling unisexual salamanders of different ploidy levels (2N vs 3N vs 4N).

The implications of these endurance disparities are ecologically and evolutionarily significant. In pond-breeding amphibians, locomotor capacity is the fundamental trait dictating the ultimate success of terrestrial dispersal (Cayuela et al., 2020; Denton et al., 2017; Joly, 2019; Semlitsch, 2008). Dispersal is the critical demographic mechanism utilized to colonize newly created habitats, escape competition in natal pools, and maintain gene flow across metapopulations (Le Galliard et al., 2003; Marsh and Trenham, 2001). While dispersal success is mediated by cognitive traits such as exploration, boldness, and risk-taking (Bredeweg et al., 2019), locomotor endurance defines the physiological boundary of dispersal capacity—the maximum distance an individual can travel before exhaustion and desiccation (Bennett et al., 1999; Hillman et al., 2014). The significantly reduced endurance of unisexual *Ambystoma* is likely to constrain their maximum lifetime dispersal capacity, placing unisexual salamanders at a competitive disadvantage compared to more mobile sexual species. Conversely, the superior endurance observed in sexual species equip them to be more efficient colonizers and potentially results in less energy intensive yearly migrations to and from breeding wetlands.

Differences in performance between *A. laterale* and *A. jeffersonianum* can potentially be explained by their specific morphology and natural history. *Ambystoma jeffersonianum* demonstrated the highest overall endurance capacity, and this relatively large species displays a strong affinity for upland, well-drained, unglaciated deciduous forests (Petranka, 1998). The topographical complexity of these upland habitats may place a strong evolutionary premium on locomotor efficiency (Bennett et al., 1989). In contrast, *A. laterale* exhibits a more compact, stockier morphology with noticeably shorter limbs and digits (Klemens, 1993; Petranka, 1998). This species is associated with flatter, glaciated lowland environments and swamps, where the distances between terrestrial refugia and breeding sites are often shorter, and the topographical resistance is inherently lower. Differences between dispersal capability between unisexual and sexual salamanders is likely to play some role in how unisexual salamanders and their sexual hosts maintain coexistence despite their reproductive overlap and presumed competition (Denton et al., 2017; Greenwald et al., 2016; Mills et al., 2020). Paradoxically, a dispersal asymmetry may facilitate the persistence of unisexual populations: if sexual hosts disperse more broadly and maintain higher connectivity between populations, they may prevent host depletion and ensure that unisexuals maintain access to sperm donors without driving local sexual populations extinct (Kerr et al., 2006; Kokko et al., 2008). Using the results from this study can be a helpful parameter to include in these models as scientists work to gain long term data on unisexual salamander populations and understand these unisexual-sexual dynamics.

## Acknowledgements

We wish to acknowledge the many individuals responsible for acquiring permitting and collecting animals for this work, including but not limited to Greg Burns, Cy Mott, Andrea Drayer, Josh Dyer, Jill Leonard, Brittany Mosher, Dennis Quinn, Matt Dykstra, Al and Lauren Blyth, John Iverson, Dick Durtsche, Stephanie Schelble, Kirsten Nicholson, Bryce Wade, Eli Bieri, Allegra Mitchell, Stephen Richter, Tim Matson, Roberta Muehlheim, Greg Lipps, Mike Benard, and Bobby Arnold. We acknowledge the contributions of M.W. Itgen and B. Arnold for their assistance during experiments.

## Author Contributions

BM and RDD conceptualized this project, and data was collected by BM, JC, and GB. Statistical analyses were performed by JC and GB. BM wrote the original draft of this manuscript, RDD reviewed and edited, and all authors reviewed prior to submission. RDD supervised all stages of the research.

## Data Availability Statement

All data and code used to conduct analyses are included as a supplement for review and at a private Dryad Digital Repository (link: http://datadryad.org/share/LINK_NOT_FOR_PUBLICATION/sZqMVywyNQuAudu6YBMYhsvnC74r_FzbuGFaA9CEM-w). Upon acceptance for publication, the repository will be made public.

## Ethics Statement

The animals used in this study were obtained following local state wildlife regulations as managed by collectors in the acknowledgements and housed under Marian University IACUC #RD2021-2.

